# Indications of an *Achaea* sp. caterpillar outbreak disrupting fruiting of an ectomycorrhizal tropical tree in Central African rainforest

**DOI:** 10.1101/2022.10.18.512753

**Authors:** Julian M. Norghauer, David M. Newbery, Godlove A. Neba

## Abstract

**Background and aims:** Where one or several tree species come to dominate patches of tropical forest, as many masting ectomycorrhizal legumes do in central Africa, ecological theory predicts they may be prone to herbivory which might alter their reproductive output. This was indirectly investigated in lowland rainforest in Cameroon for *Tetraberlinia korupensis*, whose crowns were attacked in 2008 by an outbreaking black caterpillar—identified as an *Achaea* sp., probably *A. catocaloides—*in Korup National Park.

**Material and methods:** Field-collected data on tree-level seed and fruit (pod) production of *T. korupensis* in its 2008 masting event were compared with that of its two co-dominant neighbours (*T. bifoliolata, Microberlinia bisulcata*) whose populations masted in 2007 (and 2010). To do this, bivariate regression models (linear, polynomial, ZiG [zero-inflated gamma model]), contingency table analysis, and non-parametric measures of dispersion were used.

**Key results:** Assuming *T. korupensis* is prone to *Achaea* caterpillar attacks, empirical data support the hypothesized lower proportion of adults participating in its masting (54% in 2008) than for either masting population of *M. bisulcata* (98% in 2007, 89% in 2010) or *T. bifoliolata* (96% in 2007, 78% in 2010). These fruiting *T. korupensis* trees were about one-third larger in stem diameter than conspecific non-fruiters and produced as many pods and seeds per capita as *T. bifoliolata*. But regressions only modestly support the hypothesis that the positive tree size–fecundity relationship for *T. korupensis* was weaker (i.e., lower adj. *R^2^*) than for *M. bisulcata* (whose leaves are morphologically similar) or *T. bifoliolata*, with mostly similar dispersion about the median among these species.

**Conclusion:** Altogether, the findings suggest a role for tolerance in nutrient-poor forests. It is postulated that instead of conferring resistance to herbivores, the ectomycorrhizas associated with these trees may enable them to more quickly recover from potential yet unpredictable insect outbreaks.

## INTRODUCTION

In tropical forests, certain tree species are capable of dominating the canopy at neighbourhood scales and are usually ectomycorrhizal (EM) and mast fruiting/seeding (Connell and Lowman 1989, Newbery et al. 1998, 2013, Torti et al. 2001, Peh et al. 2011, Corrales et al. 2018). The linkage between these two traits was articulated in Newbery’s (2005) commentary on Henkel et al. (2005). Yet two aspects of their biology are still poorly known relative to temperate forests. The first is the prevalence of insect herbivore outbreaks in the canopy (Nascimento and Proctor 1994, Maisels 2004, Dyer et al. 2012), and the second is how intraspecific variation in seed (or fruit, capsule, pod) production varies with individual tree size (e.g., Norghauer and Newbery 2015). Both require empirical investigation to address whether and how the former impairs the latter (Rockwood 1973, Wong et al. 1990). This could lead to a better understanding of the population dynamics, reproductive phenology, and species coexistence of trees (Dyer et al. 2012), as well as improving forest management and conservation efforts (Nair 2007), especially under a changing climate (Pureswaran et al. 2018).

Whereas insect outbreaks in temperate forests are well-studied phenomena (Kulman 1971), notably for tree-defoliating pests (Johns et al. 2016), the same cannot be said for tropical forests outside of plantations (Gray 1972, Nair 2007). For most of the last century it was wrongly believed that outbreaks were rare or absent in these communities, a view no longer tenable given the review by Dyer et al. (2012). They argued that insect outbreaks in the tropics might be as frequent and severe as in temperate zones, being more likely to happen where hosts are most concentrated (aggregated or clumped), in line with ecological theory predicting high damage levels to plant monocultures from herbivory (the ‘resource concentration hypothesis’ of Root 1973) and related modelling of host-plant dynamics (Stephens and Myers 2012). Accordingly, being akin to them in natural settings, dominant species with EM and mast fruiting and limited dispersal (clumped adults or groves) could be prime candidates prone to such outbreaks, though these events are likely cryptic and harder to notice in tropical forests because of lianas and denser vegetation. Although its crown foliage has never been surveyed for outbreaks, saplings of the monodominant *Gilbertiodendron dewevrei* tree in the Congo incur greater herbivory to their leaves from insects than do nearby mixed-forest species, suggesting that EM does not confer resistance as originally theorized (Gross et al. 2000).

In tropical forests, little effort has been devoted to *direct* counts of seeds (or fleshy fruits, capsules, or pods) at the tree level—not the stand or population level—across sufficient numbers of individuals (i.e., spanning the species’ full range of stem diameters). To better understand tree evolutionary ecology and size–fecundity relationships vis-à-vis insect outbreaks, especially for those with masting populations (synchronized supra-annual seed production; Kelly and Sork 2002), data are needed *from individual trees from relatively intact forests—*not a new argument or plea (Janzen 1978, Herrera et al. 1998, Koenig et al. 2003). Such investigations remain limited, however (e.g., Kainer et al. 2007, Klimas et al. 2012, Norghauer and Newbery 2015, Minor and Kobe 2019), and although seed production generally increases with plant size, this size–fecundity relationship generally declines in trees due to senescence and physiological decline (Qiu et al. 2021).

Ideally, both aspects above would be tackled and bridged simultaneously by monitoring tree species’ adults over many years. When that is not possible, opportunistic observations of an insect herbivore’s presence and activity are still useful when coupled with empirical data on tree–level fecundity. To that end, this study focused on *Tetraberlinia korupensis* in southwest Cameroon. The aims were (1) to briefly note the occurrence of an outbreaking caterpillar (*Achaea* sp.) in crowns of this tree species in 2008; (2) to test hypotheses H_M_, H_S1_, and H_S2_ (as set out below) concerning the variation in reproductive output around that phenomenon among co-occurring dominant species; and (3) to consider further the impacts of insect outbreaks on tropical tree reproductive phenology and fruiting–size relationships for ectomycorrhizal species. Due to the remoteness of the study site, a long time series of recorded insect feeding and tree responses was not achievable, but nevertheless the study serves as a means to a more refined understanding of tree growth and allocation, leading to the postulate of ectomycorrhizal-mediated tolerance to insect attack.

We expected to find that the general size–fecundity relationship of *T. korupensis* is more variable (i.e., harder to predict individual fecundity on the basis of size alone) in comparison with one or more co-dominant masting species either free of or less prone to outbreaks (H_M_, main hypothesis). If supported, this would be consistent with differential damage incurred across the *T. korupensis* population and, perhaps more likely, differential responses of individuals to herbivory. Both processes will probably be modulated by host tree size and vigour (Price 1991), intraspecific variation in light availability and leaf nutrients (Dudt and Shure 1994), and the palatability and density of neighbouring trees not targeted by the herbivore (Barazza et al. 2006). Implicit in this reasoning is that past herbivore outbreaks have occurred that could affect reproduction tightly coupled to host tree size. We also anticipated that, compared with the same co-dominant species, a lower proportion of adult-sized trees of *T. korupensis* would participate (i.e., actual “fruiters”, those producing pods containing seeds) in a masting event (H_S1_, subsidiary hypothesis 1) and the dispersion (*CV*, coefficient of variation) in tree-level fecundity of these non-zero fruiters would be higher (HS2, subsidiary hypothesis 2). Support for the former would be consistent with insufficient resources available (after replacing eaten vegetative tissues) to produce seeds from recurring bouts of severe defoliation. The latter would particularly apply to a mast fruiting strategy that operates via an internal resource threshold that must be crossed to fruit (Kelly and Sork 2002, Newbery et al. 2006a). Furthermore, for masting to be effective in normal years, most individuals would each be contributing many seeds at each event, and more uniformly.

## METHODS AND MATERIALS

### Study site and species

Korup National Park (at 50–150 m a.s.l.; 5°10’N, 8°50’E) preserves Atlantic lowland rainforest of southwest Cameroon on nutrient-poor sandy and acidic soils, which served as a crucial vegetation refuge during the last Ice Age (Newbery et al. 1988, 1997, 1998; van der Burgt et al. 2021). There is no evidence of fire usage or commercial logging in Korup. In its southern part lies the 82.5-ha permanent ‘P-plot’, set up in 1991, whose canopy is dominated by three (Leguminosae or Fabaceae) ectomycorrhizal masting trees (subfamily: Detarioideae; LPWG, 2017): *Microberlina bisulcata* A. Chev., *T. bifoliolata* (Harms) Haumann, and *T. korupensis* Wieringa (formerly *T. moreliana* Aubr.) (Newbery et al. 1998, 2006a, 2013). Full details of the background and set-up of the P-plot are to be found in these last-cited papers. Seeds of all three species are discoid-shaped (Supplementary file: Fig. S1) and ejected from pods atop crowns on sunny days in the wet season (late June to early September).

That *T. korupensis* is a masting tree species is supported by several lines of evidence. Firstly, in 2007 and 2010, respectively, few if any *T. korupensis* seeds were caught (J. M. Norghauer, pers. obs.) in 580 traps by Norghauer and Newbery (2015; numbers not reported there), and extensive searching across 82.5 ha in December 2007 found newly established *T. korupensis* seedlings in just 10 canopy gap areas, these located in the 1/3-westernmost portion of the P-plot where adult densities are highest (Newbery et al. 2013). This contrasts starkly with 2008, when *T. korupensis* seeded heavily, and its seedling recruitment in terms of density (N per ha) was near double that recorded in 1995 when all three species had flowered and seeded heavily (Newbery et al. 1998), and it exceeded that of *T. bifoliolata* in the latter’s 2007- and 2010-confirmed mast seeding events (Supplementary file: Table S1). Furthermore, new seedlings of *T. korupensis* in 2008 were slightly more and less frequently present in plots than those of *T. bifoliolata* in 2007 and 2010, respectively (Table S1). Thirdly, over a 10-year period (1988–1997), collectively, field workers noted substantial seeding and seedling recruitment in just two or at most three years (if 1997 is included, Table S1; Newbery et al. 1998, Newbery et al. 2006b), this consistent with masting behaviour (i.e., intervals of at least one year with very low or negligible reproduction).

### Tree-level fecundity measurements

In late December 2008/early January 2009, all *T. korupensis* trees in the eastern-most 25-ha part of the P-plot were visited—plus those in a 50-m buffer on its N, S, and W sides, and their reproductive output from mast-fruiting in August–September 2008 quantified (i.e., fallen pod valves under the crown) (Fig. S1). At every tree ≥ 10-cm DBH (diameter at breast height; these measured last in 2005; Newbery et al. 2013), its crown edges were measured along six bearings (60° increments) to the nearest 0.5 m, aided by a right-angle prism. Then, four 1-m radius circular ground plots (each 3.14 m^2^ in area) were set up *midway* along the tree crown extensions on the four cardinal bearings (N, E, S, W), this positioning guarding against sampling pod valves possibly originating from a large neighbouring conspecific fruiter (an issue that arose in Green and Newbery 2002). In each plot, all found pod valves were hand-counted (two valves = one pod), and their respective seed scars also counted (Fig. S1). This sampling methodology was exactly the same as that used for *M. bisulcata* and *T. bifoliolata* populations in their 2007 joint masting events (and also in 2010; their data collected and reported in Norghauer and Newbery 2015). An individual tree’s horizontal crown cross-sectional area (m^2^) was calculated as the sum of the six triangles forming a polygon. Average pod count per m^2^ across the four plots (i.e., first dividing by ground plot area) was multiplied by its crown area (m^2^), to obtain total number of pods per tree, and this tally multiplied by the average scar count per pod to obtain total number of seeds per tree. Both variables served as estimates of tree-level reproductive output (i.e., fecundity).

### Outbreaking caterpillar and size range of *T. korupensis* trees attacked

An insect outbreak of a black morphotype caterpillar (Fig. 1) on *T. korupensis* trees occurred in mid to late December 2008. Later, on 18-19 March 2009, 19 trees opportunistically recorded as attacked (frass present or caterpillars seen) in December 2008 were measured for stem girth—14 in the P-plot’s eastern 25-ha; four in its western part, plus another one midway. The range in DBH was 69 to 140 cm. For the 14 attacked *T. korupensis* trees for which crowns were measured, their areas ranged from 76 to 616 m^2^. Of these, three had failed to produce any pods/seeds and were clustered in the P-plot, consistent with an insect outbreak having an epicenter (Nair 2007).

**Fig. 1.**
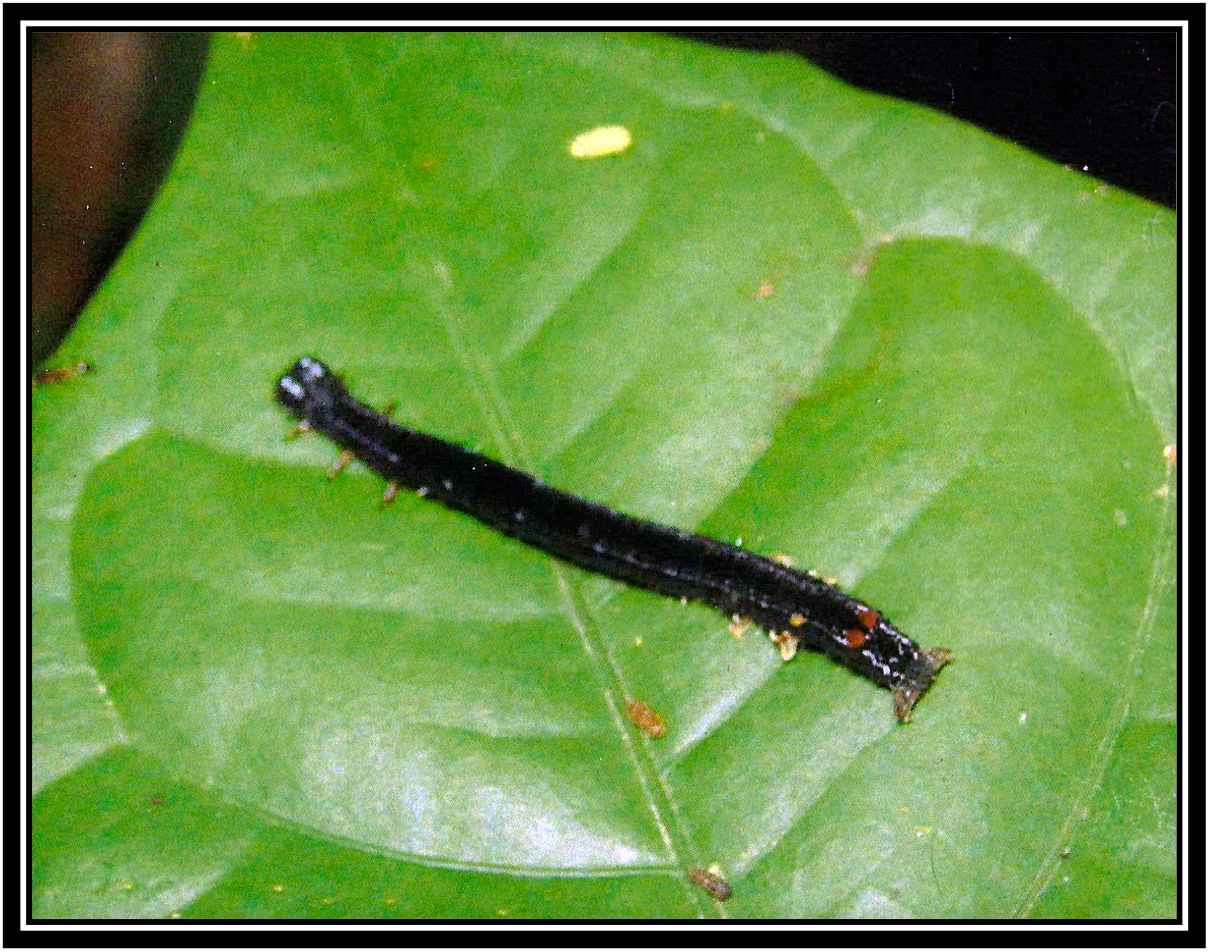
Photograph of the *Achaea* caterpillar (early to mid instar, dorsal view)—here seen resting on an unidentified heterospecific plant in the forest understorey—that was active in the crowns of *Tetraberlinia korupensis* trees in Korup National Park, Cameroon. Photo credit: Godlove A. Neba.

### Data processing and statistical analyses

Accordingly, a rounded-down lower limit to crown size was taken at 75 m^2^ for the inclusion of trees in the statistical analyses. This constraint was required to compare tree species on a common crown area basis, because female egg-laying moths of the caterpillar likely rely on a minimal density or area of flushing foliage in the forest canopy as a foraging cue for a host tree of suitable size (Nair 2007; e.g., Haugaasen 2003); hence, it was assumed that the caterpillar did not typically attack trees with a crown area ≤ 75 m^2^. An upper threshold was not imposed, however, because this would have excluded many larger *M. bisulcata* trees whose fecundity might inherently be more variable due physiological decline as well as crown thinning (Norghauer, pers. obs.; Newbery et al. 2006a) and whose leaf flushes might be more apparent or attractive to herbivores. The largest crown of an *M. bisulcata* tree (1458 m^2^) was almost three times that of *T. bifoliolata* (470 m^2^) or *T. korupensis* (616 m^2^). The lower 75-m^2^ cut-off was applied to all trees above the size onset of maturity (SOM), that is 41.9 cm DBH for *M. bisulcata* and 25.4 cm DBH for *T. bifoliolata* (Norghauer and Newbery 2015), with the latter presumed to be the same for the similarly statured *T. korupensis* (Newbery et al. 2013). Further details of the sample selection are in the Supplementary file: Methods S1.

#### Size–fecundity relationships of the three species

These were first explored by fitting generalized additive models (GAMs), using the ‘mgcv’ package (Wood 2020) under its default settings in R v4.1.3 (R Core Team 2022). A GAM does not presuppose a specific functional relationship and can objectively find a best-fitting smoother—a thin plate regression spline—by minimizing the criterion of GCV (generalized cross-validation; Wood 2004). For the ln-transformed counts of seeds (and pods), these GAMs clearly revealed linear relationships for both *Tetraberlinia* species but a curvilinear (hump-shaped) one for *M. bisulcata* (data not shown). Accordingly, to test HM, simple linear and polynomial regression models were used, for which the proportion of variation explained in seed (or pod) production by tree stem size (DBH) was compared using the adjusted *R^2^*-value. This goodness-of-fit measure takes into account the sample size *N* (Agresti 2015; p. 56), thus enabling a fairer (less biased) comparison among species. Crown area could not be used as the *X* variable, as this could lead to spuriously inflated *R^2^*-values (Brett 2004). It was not mathematically independent of pod counts because crown area was used in their calculation, and thus seed production (the *Y* variable). For the six *T. korupensis* fruiting trees in the 50-m buffer, DBHs had to be interpolated from their measured crown areas, using the species-specific fitted linear relationship: log_10_(DBH) = 0.6973 + 0.5128(log_10_[crown area]), *P* < 0.001, *R^2^* = 0.764, *F*_1, 137_ = 474.7; Fig. S2). All these regressions were fitted in Genstat v16.2 software (VSN International Ltd, Hemel Hempstead, UK).

Lastly, to take into account the relatively many *T. korupensis* trees (compared with only two each of *M. bisulcata* and *T. bifoliolata*) that produced no pods or seeds, a zero-inflated gamma (ZiG) model was fitted to the complete data (i.e., zero and positive-count outputs together) following the approach of Zuur and Ieno (2016: pp. 113–132). This hurdle type of model combines a fitted binomial (Bernoulli) generalized linear model (GLM) on presence/absence of output with a separately fitted gamma GLM, with the log-link, on just the positive output values (seed or pod counts per tree). Goodness-of-fit was assessed by McFadden’s (1974) pseudo-*R*^2^ value, this based on the log-likelihood ratio of the fitted to a null model (see Mittelböck and Schemper 1996, and Hosmer et al. 2013: pp. 182–186). Log-likelihoods from the binomial and gamma fits were added together for each of the fitted and null models (all implemented in R [R Core Team 2022]).

#### Participation in masting events

To test H_S1_, because of low expected cell counts < 5, permutations (n = 4999) were used for a 2 × 2 contingency table analysis that compared the proportion of trees fruiting in the *T. korupensis* population to that of *M. bisulcata* or *T. bifoliolata*. For comparative purposes, the opportunity was taken to derive the SOM for *T. korupensis*, in the same way as done earlier for its two co-dominant species in Norghauer and Newbery (2015), by fitting a modified logistic regression and calculating *Dcrit*—using eqs 3 and 5 in Thomas (1996)—but to all available fecundity data (i.e., 164 trees ≥ 10 cm DBH scored for yes [1] vs no [0] seed/pod production). For both analyses (and summary statistics) Genstat v.16.2 was used.

#### Dispersion of fruiting trees’ seed and pod production

To test H_S2_, for each population, because of their right-skewed distributions for non-zero seed and pod counts, two recognized non-normal measures of relative dispersion were considered, for which the calculated medians omitted the non-fruiting trees (i.e., zero count data). The first is based on the interquartile range (*CQV:* coefficient of quartile variation) with its 95% CI (confidence interval), obtained using the method of Bonett (2006) in R with the ‘cvcqv’ package (Beigy 2019). The second is based on the mean absolute deviation from the median, whose ratio to the median can be denoted as *CV*_MnMad_ following Ospina and Marmalejo-Ramos (2019). This is also known as the ‘coefficient of dispersion’ and was calculated using the ‘statpsych’ package in R (Bonett 2021), for which Bonett and Seier’s (2006) improved non-bootstrapping method was used to derive the 95% CIs.

## RESULTS

### Caterpillar description and putative identification

A black caterpillar (Fig. 1) attacked *T. korupensis* crowns in mid to late December 2008 (ca 2-week period), this being early in the dry season when this species (and *M. bisulcata*) annually exchanges its foliage. Caterpillars ate flushing or newly expanded leaves, moved in a looping way, were highly gregarious, and made a distinctive pitter-patter sound while chewing and defecating (falling frass hitting leaves below). They descended or fell to the understorey on silken threads, some returning on them, others quickly scrambling up tree trunks. As this crown defoliation happened *after* the *T. korupensis* population had mast fruited and dispersed its seeds in 2008, the analyses presented below assumed a history of unobserved outbreaks on *T. korupensis*. Morphologically, this small non-hairy caterpillar has three pairs of laterally protruding true legs and what seems to be only two or three usable prolegs (two ventral plus anal claspers), and a pair of pronounced red diamond-shaped knobs on its back (Fig. 1). After consulting with lepidopterists (Dino Martins and Scott Miller, pers. comm.) it was confirmed to be a noctuid moth, specifically an *Acheae* sp., most likely *A. catocaloides* Guenee (if not the similar-looking outbreaker *A. lienardi* Boisduval) in the family Eribidae, superfamily Noctuoidea (http://www.afromoths.net/species/show/33376 [De Prins and De Prins 2011-2021]), whose larvae could also be heard feeding and climbed silky threads as they defoliated many mature trees of the monodominant tree *Paraberlinea bifoliolata* Pellig. (Fabaceae) in Gabonian rainforest (Maisels 2004). Present across tropical Africa, *A. catocaloides* larvae are highly polyphagous, being not only known pests of various crops (Latham et al. 2022), including cacao in Cameroon (Dejean et al. 1991), but also commonly found on the legume forest trees *Pentaclethra macrophylla* Benth. (Eluwa 1977) and *P. eetveldeana* De Wild. & T. Durand (Latham et al. 2022), whose pinnate leaves are morphologically very similar to those of *T. korupensis* (and *M. bisuclata*).

### Tree-level fecundity (seed and pod production)

For all three species their tree-level seed production was skewed, with medians half to one-third their corresponding mean values (Table 1). Among *M. bisulcata* individuals, seed production varied at least 5000-fold, but less so in the populations of *T. bifoliolata* (≈ 350-fold) and *T. korupensis* (≈ 200-fold). Whether gauged by the mean or median, seed production was 2–3 times more for *M. bisulcata* than for *T. bifoliolata* and *T. korupensis;* however, expressed as per capita pod production, it was actually higher for *T. korupensis* than *T. bifoliolata*. There is close similarity in the tree-level fecundity (seed or pods) between *T. korupensis* and its congener.

**Table 1.**
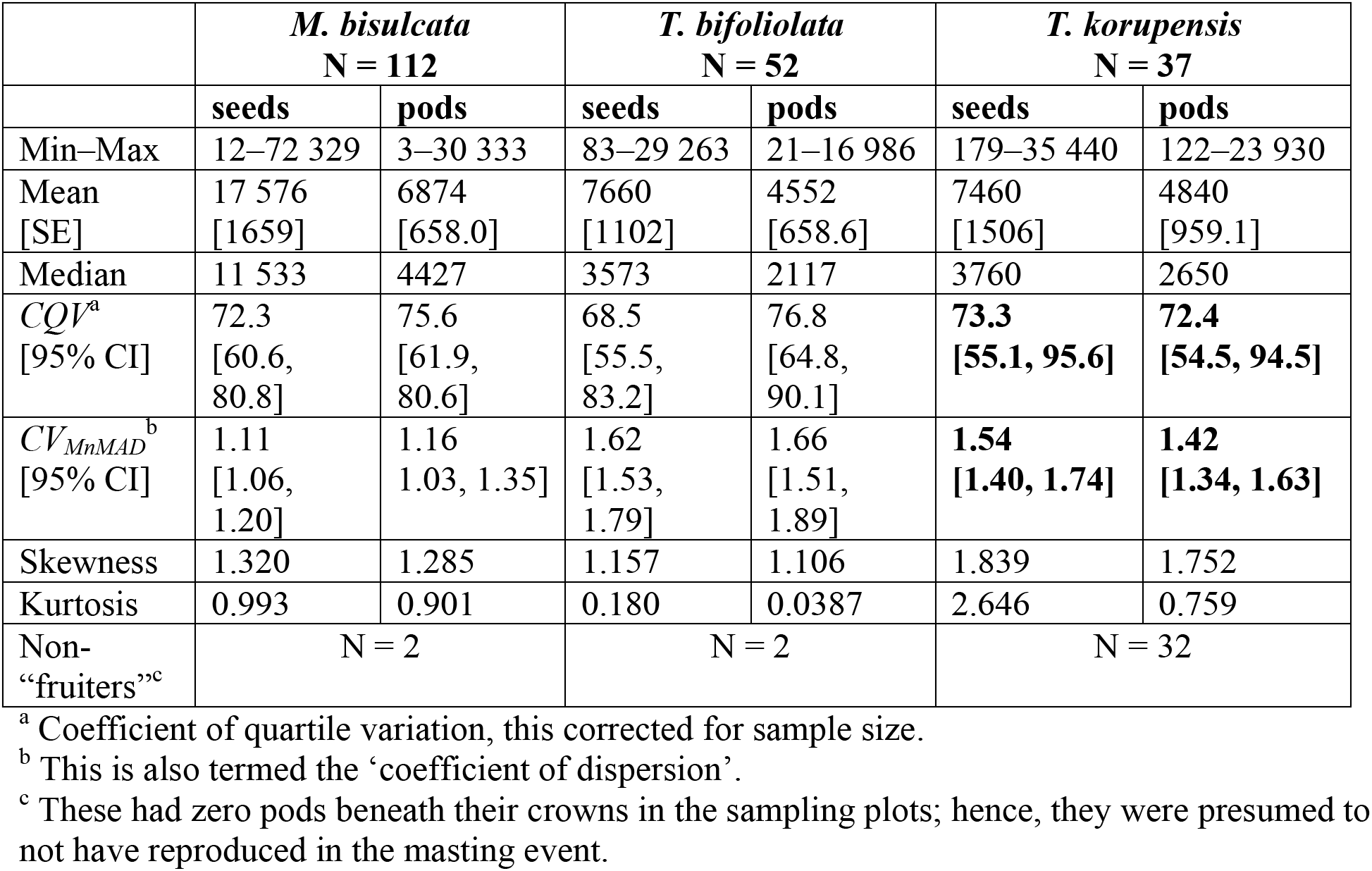
Statistics for each species’ population subsample of adult-sized trees (based on tree stem diameter, DBH) having a crown area greater than 75 m^2^ in the 2007 masting event (*Microberlinia bisulcata* [total N = 114] and *Tetraberlinia bifoliolata* [total N = 54]) and the 2008 masting of *T. korupensis* (total N = 69) in Korup National Park, Cameroon.

|  | *M. bisulcata*<br>N = 112 |  | *T. bifoliolata*<br>N = 52 |  | *T. korupensis*<br>N = 37 |  |
| --- | --- | --- | --- | --- | --- | --- |
|  | seeds | Pods | seeds | Pods | seeds | Pods |
| Min–Max | 12–72 329 | 3–30 333 | 83–29 263 | 21–16 986 | 179–35 440 | 122–23 930 |
| Mean<br>[SE] | 17 576<br>[1659] | 6874<br>[658.0] | 7660<br>[1102] | 4552<br>[658.6] | 7460<br>[1506] | 4840<br>[959.1] |
| Median | 11 533 | 4427 | 3573 | 2117 | 3760 | 2650 |
| *CQV*<sup>a</sup><br>[95% CI] | 72.3<br>[60.6, 80.8] | 75.6<br>[61.9, 80.6] | 68.5<br>[55.5, 83.2] | 76.8<br>[64.8, 90.1] | **73.3**<br>**[55.1, 95.6]** | **72.4**<br>**[54.5, 94.5]** |
| *CV<sub>MnMAD</sub>*<sup>b</sup><br>[95% CI] | 1.11<br>[1.06, 1.20] | 1.16<br>[1.03, 1.35] | 1.62<br>[1.53, 1.79] | 1.66<br>[1.51, 1.89] | **1.54**<br>**[1.40, 1.74]** | **1.42**<br>**[1.34, 1.63]** |
| Skewness | 1.320 | 1.285 | 1.157 | 1.106 | 1.839 | 1.752 |
| Kurtosis | 0.993 | 0.901 | 0.180 | 0.0387 | 2.646 | 0.759 |
| Non-<br>“fruiters”<sup>c</sup> | N = 2 |  | N = 2 |  | N = 32 |  |

<sup>a</sup> Coefficient of quartile variation, this corrected for sample size.
<sup>b</sup> This is also termed the ‘coefficient of dispersion’.
<sup>c</sup> These had zero pods beneath their crowns in the sampling plots; hence, they were presumed to not have reproduced in the masting event.

#### Participation in masting events

In the *T. korupensis* population only 53.6% of the adult-sized trees ≥ 75 m^2^ in crown area fruited in 2008 (Table 1), which differed significantly from nearly all (98.0%) for *M. bisulcata* (likelihood χ^2^ = 60.3, *P* < 0.001) and 96.3% for *T. bifoliolata* in their 2007 masting event (likelihood χ^2^ = 32.6, *P* < 0.001). The null hypothesis of equal proportions was therefore rejected in each case (hypothesis H_S1_). Those *T. korupensis* trees that did seed were at least one-third larger in stem size (DBH) than non-fruiters (means ± SD: 87.3 ± 24.1 vs 62.0 ± 22.5 cm, *t* = −4.49, *P* < 0.001, d.f. = 67). If instead all adults irrespective of their crown areas are included, 42.3% (44/104) of *T. korupensis* trees reproduced; and even when the SOM of this species is increased to 35 cm DBH, just 51.2% reproduced (43/84). But doing the same (relaxing the 75-m^2^ cut-off) for *T. bifoliolata* decreased its proportion of masting participants to 73.0% (73/100). For the empirically derived SOM of *T. korupensis* based on all available data in 2008, the *D_crit_* corresponding to the modal stem diameter (DBH) of first reproductive output was 53.2 cm (modified logistic regression parameters: *a* = −12.35, *b* = 2.92, df = 2, 162), this 11 and 28 cm greater than that of *M. bisulcata* and *T. bifoliolata* in 2007, respectively.

#### Dispersion of fruiting trees’ seed and pod production

For the dispersion of untransformed tree-level fecundity estimates, among the three species (hypothesis HS2), the *T. korupensis* population did have the highest *CQV*-value for seed production, but only by a very small margin; for pod production its *CQV* was lowest (Table 1). Actually, within species, *CQV* was quite similar between seed and pod counts for *M. bisulcata* and *T. korupensis*, and least so for *T. bifoliolata*. Furthermore, 95% CIs for the *CQVs* overlapped among the three species. By contrast, the two congeners had similar values for *CV*MnMAD (seeds and pods), these 22%–43% higher than those of *M. bisulcata*. In this last case, however, significantly greater dispersion characterized the seed fecundity of *T. korupensis* (i.e., 95% CIs non-overlapping with those of *M. bisulcata*), and nearly so for pods (Table 1).

#### Size–fecundity relationships of the three species

As Fig. 2 shows, stem size (DBH) accounted for just over 2.5-fold more of the variation in seed production of *M. bisulcata* (*F*_2, 109_ = 38.5, *P* < 0.001; Fig. 2A) than *T. korupensis* (*F*_1, 35_ = 7.3, *P* = 0.010; Fig. 2C). For the *T. bifoliolata* population, nearly 25% of the variation in fecundity could be explained by stem size (*F*_1, 50_ = 17.5, *P* < 0.001; Fig. 2B) despite two outlying values, this being 1.66 times that of *T. korupensis* (Fig. 2C). A similar weaker ability of individual tree size to predict seed production in the *T. korupensis* population (hypothesis HM) was found for pods (adj. *R^2^* = 0.169, *F*_1, 35_ = 8.31, *P* = 0.007;) vis-à-vis *M. bisulcata* (adj. *R^2^* = 0.420, *F*_2, 109_ = 41.2, *P* < 0.001) and *T. bifoliolata* (adj. *R^2^* = 0.261, *F*_1, 50_ = 19.0, *P* < 0.001).

**Fig. 2.**
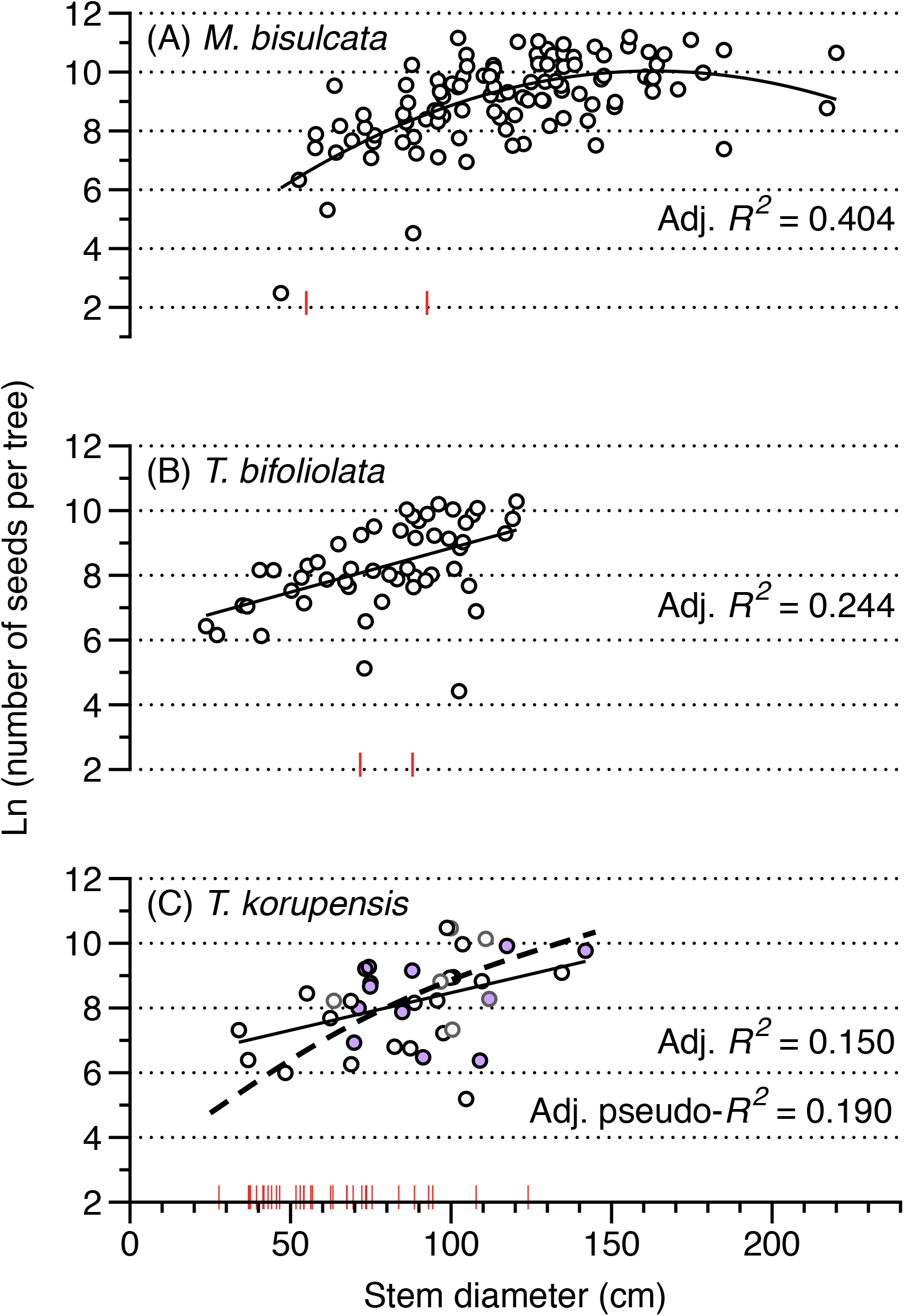
Scatterplots of individual reproductive output (seeds) as a function of stem diameter for three canopy-dominant masting species—*Microberlinia bisulcata, Tetraberlinia bifoliolata*, and *T. korupensis*—in Korup National Park, Cameroon. Red vertical tick marks along the *X*-axis indicate the size of non-fruiting trees (no seeds or pods produced) in each population. In (**C**), the light-gray circles are six trees whose stem diameter was interpolated from their crown areas (Fig. S2); purple-filled circles are those trees confirmed as attacked by the *Achaea* caterpillar. Another three scored as attacked did not fruit (red ticks; DBHs: 67.5, 73.7, and 124.0 cm). The dashed curved line is the fitted ZiG (zero-inflated gamma) regression to the 69 *T. korupensis* trees (i.e., fruiters plus non-fruiters). Because the scatterplots for pods closely resembled these for seeds they are not shown for brevity. Samples sizes for fruiting trees only are (**A**) N = 72, (**B**) N = 52, and (**C**) N = 37.

For the ZiG model of *T. korupensis*, the binomial GLM fit of 0/1 reproductive output as a function of stem diameter was highly significant (deviance ratio = 15.34, df = 1, 67; *P =* 0.000213; estimated dispersion = 1.16). The gamma GLM fits, for both seeds and pods positive counts on diameter, were also highly significant (*t* = 4.047 and 4.310 respectively, df = 1, 35; *P* = 0.000272 and 0.000126), and more significant than the linear fit of log-counts on diameter. The estimated ZiG curve for seed counts is added to Fig. 2C; with a pseudo-*R^2^* at 19.0%, this is slightly better than for the linear fit of natural log-counts and closer to the *R^2^*-value for *T. bifoliolata*, yet still much lower than that for *M. bisulcata*. The corresponding fit for pod counts had a pseudo-*R^2^*-value of 19.7%. Although absolute variability in fruiting increased by including the zero-producers in the full tree sample (N = 69), the fit with the ZiG model using them improved over the GLM without them (N = 37) partly because the sample size nearly doubled.

When the assumption that only trees > 75 m^2^ are attacked by the *Achaea* caterpillar was relaxed, thus including other presumed adults, a different picture emerged. There was a much less variable relationship for *T. korupensis*, with adj. *R^2^*-values of 0.279 and 0.297 for seeds and pods (respectively, *F*_1, 42_ = 17.6 and 19.2, *P* < 0.001) that were a little higher than those of *T. bifoliolata* which instead required a polynomial fit (seeds, adj. *R^2^* = 0.239, *F*_2, 69_ = 12.1, *P* < 0.001, quadratic term: *P* = 0.01; pods, adj. *R^2^* = 0.217, *F*_2, 69_ = 10.8, *P* < 0.001, quadratic term: *P* = 0.01). However, with just one tree added to its sample, for *M. bisulcata*, the corresponding adj. *R^2^*-values increased only marginally (adj. *R^2^* = 0.425 and 0.440, *F*_2, 110_ = 42.3 and 45.1, *P* < 0.001), yet still greater than those of *T. korupensis*.

### Effect of a climate difference between years?

The analyses were unavoidably confounded with any differing climatic conditions between 2007 and 2008 masting years. The preceding dry season was 35 days longer in 2007 than 2008, it started also 35 days earlier, and it had 70% more mean daily rainfall in the latter (1.71 vs 1.01 mm); however, mean daily radiation and minimum temperature were similar (152 vs 158 W m^2^, 19°C vs 19°C) (D.M. Newbery, unpubl. data). If climate differences between years were confounding our comparisons between *T. korupensis* and *M. bisulcata* with *T. bifoliolata, R^2^-* values would likely have all been more similar had all three species been compared in a single year (say, in 2007 or in 2008). Fortunately, fecundity data for *M. bisulcata* and *T. bifoliolata* trees are available, albeit for smaller sample sizes, from their joint masting in 2010 (Norghauer and Newbery 2015). Although its preceding dry season was only 8 days longer than in 2008 with rainfall more similar at 1.42 mm (but higher minimum temperature at 22°C; radiation unchanged), it started even earlier than the one preceding the 2007 masting, in fact 43 days earlier than that preceding the masting in 2008 of *T. korupensis*. The mast fruiting events of *M. bisulcata* and *T. bifoliolata* in 2007 and 2010 are distinguished by their associated early starts, not so much the dry season’s entire duration. The 2008 masting of *T. korupensis* was associated with a comparatively much later dry season start with, nevertheless, an average duration and rainfall intensity.

Applying the same regressions to adult-sized stems with 75-m^2^ min. crown area for seeds in 2010, 35.8% and 51.8% of variation was explained by DBH for *M. bisulcata* and *T. bifoliolata* respectively (*F*_2, 69_ = 20.8 and *F*_1, 31_ = 35.3, both *P* < 0.001); for pods the corresponding values were 40.4% and 44.5% (*F*_2, 69_ = 25.0 and *F*_1, 31_ = 26.7, both *P* < 0.001). Relaxing the 75-m^2^ assumption still yielded high adj. *R^2^*-values: 0.351 and 0.495 (for seeds), 0.399 and 0.396 (for pods), respectively. Thus, when compared with 2007, the size–fecundity relationship for *M. bisulcata* in 2010 held up well. By contrast, the fit became even stronger for *T. bifoliolata*—in part because one possible outlier tree (#1376; DBH = 102.5 cm) in 2008 had since died—the relationship now being more predictable than that for *T. korupensis*. Although the masting participation was reduced to 88.8% (72/81) for *M. bisulcata* and 78.6% (33/41) for *T. bifoliolata* in 2010, this too, was still much greater than that for *T. korupensis*.

## DISCUSSION

### The *Achaea* caterpillar’s presence at Korup

If the *Achaea* caterpillar feeds principally on young leaves—those expanding or recently expanded leaves—then severe damage could ensue quickly, within 1 to 3 weeks, perhaps leading to premature leaf abscission. For example, within 1 week caterpillars defoliated young leaves in crowns of two monodominant leguminous tree species in tropical rainforests of Brazil and Gabon (Nascimento and Proctor 1994, Maisels 2004). Thus, such phenomena are quite easy to miss, and may explain why this caterpillar outbreaking on *T. korupensis* trees was not noticed at Korup before.

This study’s analysis presupposed past outbreaks had occurred. Indirect evidence for this can be gleaned from earlier work on site (Chuyong et al. 2000), finding not one but two or more spikes *similar in magnitude* in the amount of *T. korupensis* leaf litter (7–8 g/m^2^) collected from traps in each of two consecutive dry seasons (from May 1990–June 1992). The first spike is expected and corresponded to leaf exchange on the yearly shedding of crown foliage, but the subsequent spike(s) is harder to explain. It could have resulted from an outbreak after which damaged leaves were pre-maturely abscised, especially under drought stress conditions. After its normal crown change in the wet-to-dry season transition in 1990 and 1991, *T. korupensis* lost these new leaves soon after (litter-trapped in February-March 1991 and 1992, respectively), but interestingly, another shedding event occurred during the wet season’s peak (July). Moreover, among the 12 tree species studied, this pattern of successive, multiple sharp peaks was unique to *T. korupensis*. Further, that in either dry season the *M. bisulcata* population lacked a second spike in litterfall altogether (i.e., only its annual leaf exchange occurred), coupled with no caterpillars or signs of activity observed during extensive phenology monitoring of 150 trees (1995–2000; Newbery et al. 2006a) and 61 trees in 2009–2013 (plus some data from 2014; unpubl. data, D. M. Newbery and S. Njibili), suggests that this species is far less susceptible to regular outbreaks from the *Achaea* caterpillar. Delayed greening is one plausible explanation (Kursar and Coley 1992), as this leaf trait is associated with significantly lower damage to expanding and young leaves in the tropics (e.g., Numata et al. 2004, Queenborough et al. 2013).

Flushing leaves of *M. bisulcata* appear almost devoid of chlorophyll, initially emerging entirely white-pink, then turning purplish with hues of green, whereas those of both *Tetraberlinia* species are a mix of green and red (J. M. Norghauer, pers. obs.). Accordingly, while all species’ young leaves contain anthocyanins, *when expanding*, those of *M. bisulcata* are likely lacking most in the nitrogen nutritionally valued by insects (Mattson 1980). Yet the *Achaea* caterpillar was active again in a December 2011 outbreak and seen under some densely clustered *M. bisulcata* trees, and again in March 2012 in another grove c. 4.2 km due south (on Isangele Road; S. Njibile, pers. comm.). For these reasons, the 2008 outbreak of the *Achaea* sp. on *T. korupensis* was unlikely an ecological anomaly.

### Tree fecundity

There were clearly far fewer masting participants and a modestly less predictable size–fecundity relationship in *T. korupensis* than either *M. bisulcata* or *T. bifoliolata*, supporting a possible association between insect outbreaks and disrupted normal seed reproduction at the individual and population level (H_M_ and H_S1_, respectively). It is unlikely that that relationship is simply more variable because of shading by larger neighbours (e.g., *M. bisulcata*), since the same reasoning should apply to the similar-statured trees of *T. bifoliolata* and yet its size–fecundity relationship was still slightly less variable. Importantly, however, the difference was most pronounced vis-à-vis *M. bisulcata*, whose leaves (compound, pinnate) are morphologically nearly identical to those of *T. korupensis*. Assuming no pollinator limitation (Newbery et al. 1998), this extra “noise” could instead have arisen not only from differences in susceptibility to defoliation severity (i.e., reduced photosynthetic capacity) among *T. korupensis* individuals, but also how they may have differentially responded to (tolerated) it, which can depend on canopy shading (asymmetric competition), availability of stored reserves, rooting extent, and perhaps also aspects of their EM associations (identity of fungal species, their degree of colonization, and access to a shared network; Gehring and Bennet 2009). The last point is the most intriguing and least investigated in the tropics, both theoretically and empirically. Besides tree size, when trying to ecologically explain intraspecific variation in fecundity (seeds, fruits), researchers consider soil properties and nutrients, limited pollination, liana loads, and competition from neighbours (Kainer et al. 2007, Klimas et al. 2012, Minor and Kobe 2019). The new findings from the present study suggest caterpillars might also merit consideration for understanding the sizexy–fecundity relationships of tropical trees.

Unfortunately, fecundity data for the *T. korupensis* at Korup is currently available for only one masting event (in 2008, i.e., it is unreplicated). Ideally, sampling for tree-level fecundity in a year when all three dominant trees species masted together, as in 1995 (Newbery et al. 1998), would clearly be useful, as would finding and sampling another *T. korupensis* population where the *Achaea* caterpillar is *absent*, for an intraspecific comparison of its size–fecundity relationships. Either may be challenging without a regular field presence, or perhaps repeated aerial surveys (see Meng et al. 2018).

An insightful finding was also the very high estimate of *D_crit_* of 53.2 cm for the *T. korupensis* population. Inferring this as the actual SOM of *T. korupensis* would be mistaken, because that *D_crit_* exceeds the 41.9 cm (or 44.0 cm) in 2007 (or 2010) of the much larger-sized, more fecund *M. bisulcata* (Norghauer and Newbery 2015), whose adult mortality rate in the last decades has been almost nil (Newbery et al. 2013). This seems implausible in terms of life-history theory or tree physiology. Instead, the high 53.2-cm estimate of *T. korupensis* could be interpreted as evidence for a *shift in the SOM of seed (pod) production* due to lag or cumulative herbivory effects from one or more previous *Achaea* caterpillar outbreaks, likely exacerbated by suboptimal climatic conditions for acquiring the resources needed to fully reproduce. Hence, many of those *T. korupensis* zero-fruiters in Fig. 2C are probably “mature”, that is, capable of flowering, but depleted NSC (non-structural carbohydrates) stores precluded them from reaching the individual threshold needed to set seed. It is worth noting that its congener *T. bifoliolata* had ca 20-cm discrepancy in its *D_crit_* in 2007 (25.4 cm) vs 2010 (45.2 cm). The reasons for that shift are unknown, but it may have been driven more by abiotic than biotic external factors. Another implication is that using seed (pod/fruit) production alone may be unreliable for estimating *Dcrit* to infer the SOM of tree species. If presence/absence data for flowering are lacking, then at least multiple years of seed production data should be analysed (e.g., Norghauer and Newbery 2015).

What are the possible implications for forest dynamics at Korup? Firstly, if the *T. korupensis* population is not masting at its fullest potential, that is fewer seeds produced per capita (H_M_) and less trees fruiting in a masting year (H_S1_), then its ability to satiate predators of immature pods in crowns [pre-dispersal] or dispersed seeds [post-dispersal] might be impaired. The corollary prediction is reduced *T. korupensis* seedling recruitment in masting years affected by an *Achaea* caterpillar outbreak, and also vis-à-vis its two co-dominant competitors. Secondly, such outbreaks might disrupt the reproductive phenology of *T. korupensis*, precluding any benefit of jointly masting with *M. bisulcata* and *T. bifoliolata* to satiate seed (or pod) predators shared among them (Norghauer and Newbery 2011). Thirdly, repeated, severe *Achaea* outbreaks could lengthen the interval between mastings, effectively delaying them, or restricting the individuals that participate in them to larger-sized ones, if attacked trees require more years to store the prerequisite photosynthate and nutrients, notably P (Newbery et al. 2006a), or their ‘resource matching’ or ‘resource switching’ processes (Pearse et al. 2016) somehow become impaired. But it is not known for certain whether tree size *per se* (DBH or crown area) accurately reflects the size of an individual’s stored C and other nutrients, or its root system extent and levels of ECM colonization.

### Shifts in leaf nutrients’ status

The *Achaea* caterpillar’s activity occurred *following* a masting event, consistent with its presumed long-term presence at Korup. Yet, given that masting years can alter the nutrient status dynamics of trees, could the outbreak instead be a consequence of it? This might happen if depleted carbon (C) stores of NSC led to lower levels of anti-herbivore phenolic compounds in young leaves. There are two reasons why this is probably untenable. First, the crown foliage attacked by the *Achaea* caterpillar was a new leaf crop in 2008, not that concurrent with C resources allocated over the prior 7–9 months to reproductive parts (flowers, seeds, pods). Recent work on European beech and oak trees found no pronounced changes in the C concentrations of leaves even during their masting years (Nussbaumer et al. 2021), and the isotope-based evidence to date generally indicates negligible C limitation, in that NSC is not depleted by masting events (Han and Kabeya 2017). Second, there is compelling evidence that nitrogen (N) pools in trees are depleted after masting events (reviewed by Pearse et al. 2016, Han and Kabeya 2017), leading to reduced foliar N concentrations (e.g., Nussbaumer et al. 2021). This should, from an insect herbivore’s point of view (Mattson 1980), render any N-depleted leaves of *T. korupensis* less nutritionally attractive and therefore less—not more—sought after as food compared with other species’ foliage in the Korup forest canopy.

### Do herbivory and climate interact to influence masting?

Evidently, dry season climate can contribute to variability in size–fecundity relationships, but the new results on herbivory suggest that it is not likely to be the only factor explaining why the size-dependent reproduction of *T. korupensis* was the weakest among the three dominants. The two factors probably interact. Timing (start and duration) and intensity (mean daily rainfall) in the dry season drive leaf exchange, flush, growth and photosynthesis under high radiation conditions, which in turn leads to C gain and storage. This is thought to then bear on the frequency, interval length, and strength of mast fruiting in these caesalpiniaceous trees (Henkel et al. 2005, Newbery et al. 2006a). The timing of herbivore attacks, with the inevitable loss of new foliage, and possibly reduced C gain and storage, may plausibly control the strength and variability of masting at later dates. A late start to the dry season in one year, with probably a varying effect for different individual trees, would be even more variably affected by herbivores whose feeding is spatially heterogeneous. Tree physiological processes, which are primarily driven by climate variability, would become exacerbated by herbivore feeding, the latter preconditioning weaker (more fed-upon trees) to perform less well in the next dry season. In this way, climate and herbivore variability might interact to explain why—when settled into a longer-term quasi-equilibrium—the variation in reproductive output was higher for *T. korupensis* than *M. bisulcata* (and *T. bifoliolata*). A corollary to this might be that the *T. korupensis* population at Korup would be expected to mast at a longer interval (every 4 or 5 years) compared with the clearly established average interval of 3 years for *M. bisulcata*. Moreover, such a lack of masting synchrony would be expected to lead to partitioning of demands on soil nutrient resources, especially for P (Newbery et al. 1997), and yet still permit all three species to utilize the shared EM network (Newbery et al. 1997, 2000).

Perturbations to the forest caused by localized caterpillar attacks can be seen as interventions to the tight ectomycorrhizal-regulated nutrient cycling of the Korup ecosystem (Newbery et al. 1997, Chuyong et al. 2000). Long-term evolutionary adaptations to herbivory are almost certainly well in place in such an old, undisturbed, primary rainforest as Korup. They enable the trees to cope with occasional outbreaks, with the system returning to an equilibrium state within a few years.

### Insect outbreaks vis-à-vis the EM trait and dominance in tropical forests

Along with the evidence for only a modest to weakly disrupted size–fecundity relationship for *T. korupensis*, its adults that masted in 2008 produced as many seeds, and more pods, per capita than did *T. bifoliolata* in 2007. These results raise an intriguing ecological question. If an unavoidable consequence of [mono]dominance in the tropics is being prone to occasional insect outbreaks (e.g., Nascimento and Proctor 1994, Maisels 2004), then, compared with AM (arbuscular mycorrhizae) root symbiotic fungi, could the EM mutualism that often characterizes such tree species, especially masting ones—known to assist in resource acquisition (Newbery et al. 1998) and to protect hosts from metal toxicity and pathogens belowground (Corrales et al. 2018)—also enable their faster recovery from outbreaks? Instead of the EM trait conferring *resistance* to herbivory, for which empirical field evidence is still lacking (Gross et al. 2000; Torti et al. 2001: Table 4; Peh et al. 2011), it may nonetheless contribute to *tolerance* of it (compensatory growth, vegetative and/or generative). Mycorrhizae-mediated tolerance to plant enemies is theoretically possible (Bennett et al. 2006) and operates in a few model plants (Gehring and Bennett 2009, Tao et al. 2016). Moreover, this plant defence strategy could be augmented where dominant populations co-occur, as they do at Korup, from access to a presumably more fungally-diverse shared EM network for needed nutrients, P especially (Newbery et al. 1997, 2006a).

## CONCLUSION

To our knowledge, a role for EM in facilitating the tolerance of trees in nutrient-poor forests to herbivore outbreaks appears not have been put forward before in the literature. It may be an important factor controlling the maintenance of tropical forest dominance. While preliminary— just one record of the outbreaking *Achaea* caterpillar, probably *A. catocaloides—*this study nevertheless made use of extensive tree-level fecundity data (201 fruiting individuals) of three masting legume species across 25 ha of rainforest. Our findings suggest such an ecological role warrants further detailed investigation.

## ACKNOWLEDGEMENTS

The authors thank Albert Kembou and Pascal Ndongmo, who served as Conservator of Korup National Park during the field observations and data collection; the Ministries of Forestry and Wildlife (MINFOF) and Scientific Research and Innovation (MINRESI) in Cameroon for research permission; George B. Chuyong (University of Buea) for local coordination; and Sylvanos Njibili (late) and Charles Ohka for their excellent help in collecting trees’ crown area and fecundity data in the field. We are very grateful to Dr Dino J. Martins (Princeton University/Mpala Research Centre) and the Smithsonian Institution (Washington, D.C.) for assistance with the caterpillar’s taxonomic identification and sharing key articles on *Achaea*.

## Availability of data and material

The datasets generated during and/or analysed during the current study are available from the corresponding author upon reasonable request. The data for Newbery et al. (2013) is archived at: http://dx.doi.org/10.5061/dryad.t85n3.

## Authors’ contributions

JMN and DMN conceived the study and interpreted the results. JMN wrote the first draft and most of the manuscript, acquired the additional fecundity and crown area measurements from the *T. korupensis* population, analysed the data, and prepared the tables and figures; DMN commented, and wrote parts of the paper and fitted the ZiG model. GAN photographed the outbreaking *Achaea* caterpillar in the field and provided natural history information.

## Supplementary file

**Fig. S1.**
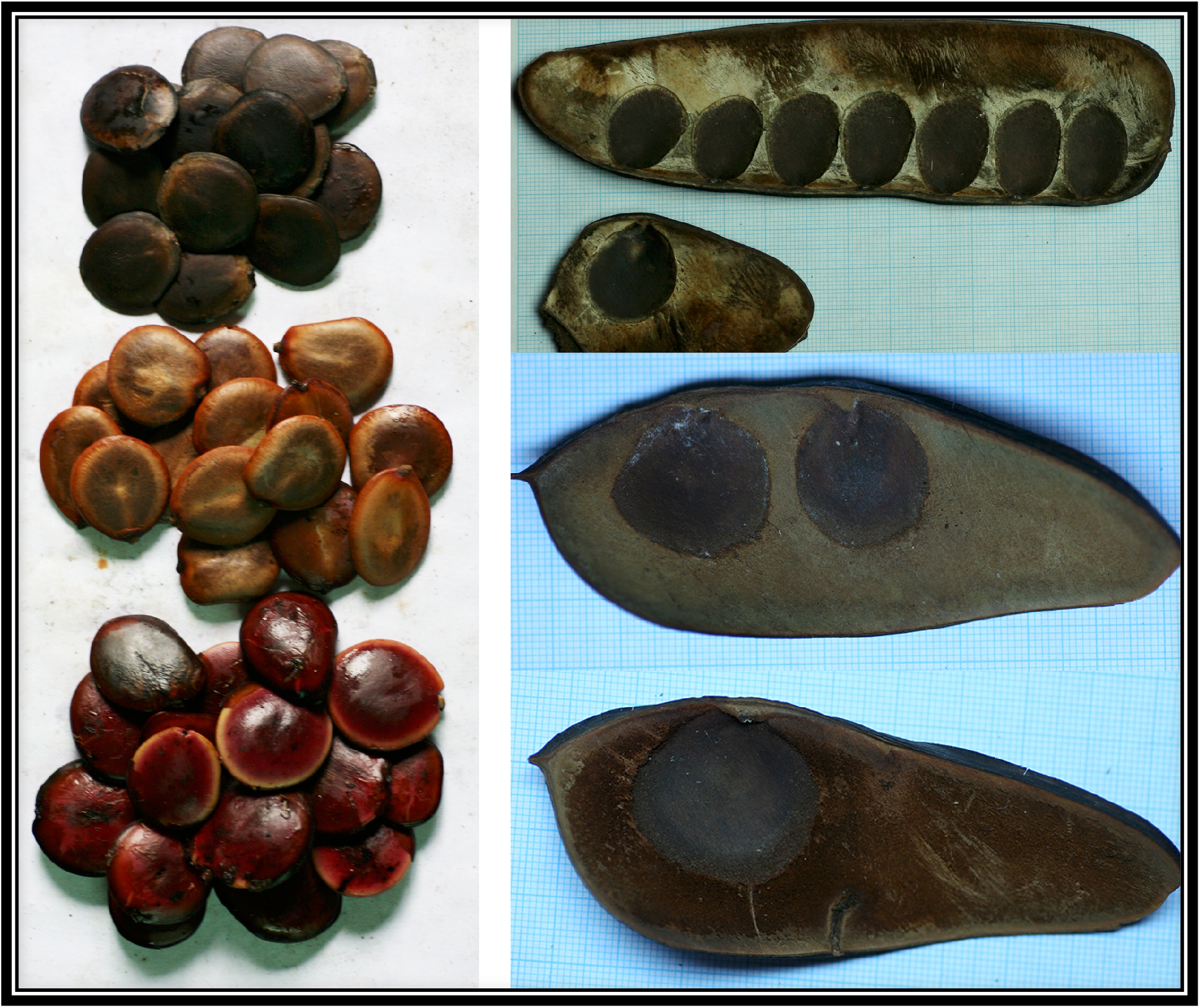
The discoid seeds (left) and pods (right) of the three co-dominant canopy tree species at Korup. From top to bottom, in order of increasing seed mass (Green and Newbery 2001): *M. bisulcata, T. bifoliolata*, and *T. korupensis*. Whereas the pods of *M. bisulcata* can house 1–7 discoid seeds, those of *T. bifoliolata* typically have one or two and occasionally three seeds, while for *T. korupensis*, which produces the largest seeds, the pod usually contains one seed only (never more than two). Dark-bordered squares = 1 cm^2^. Green JJ, Newbery DM (2001) Light and seed size affect establishment of grove-forming ectomycorrhizal rain forest tree species. New Phytologist 151:271-289

### Methods S1. Further details about the sample selection

Applying the 75-m^2^ to trees above the SOMs excluded one *M. bisulcata* (DBH = 53.8 cm, fruited in 2007), 46 *T. bifoliolata* (DBHs = 25.5–125.6 cm, of which 21 fruited in 2007), and 35 *T. korupensis* trees (DBHs = 25.6–93.0 cm, of which seven fruited). In the subsetted *T. korupensis* sample every fruiting tree was > 33 cm in DBH: one small tree (DBH = 16.1 cm, crown area = 24 m^2^) did manage to fruit in 2008, however; but no others ≤ 33 cm DBH did. Likewise, five *T. bifoliolata* trees ≤ 25.4 cm DBH fruited in 2007 (the smallest with DBH = 17.2 cm, crown area = 14 m^2^). One of these, with a crown area of 132 m^2^, but whose DBH was 23.7 cm in 2005, was added to the *T. bifoliolata* population sample because, with 2 years of growth elapsed, it would have been closer to the SOM. Although one *M. bisulcata* ‘sub-adult’ did have an 88-m^2^ crown, its 34.8-cm DBH was too far below the SOM for inclusion; another excluded tree did fruit in 2008, but its crown area was 50 m^2^ and its DBH was 37.0 cm.

**Fig. S2.**
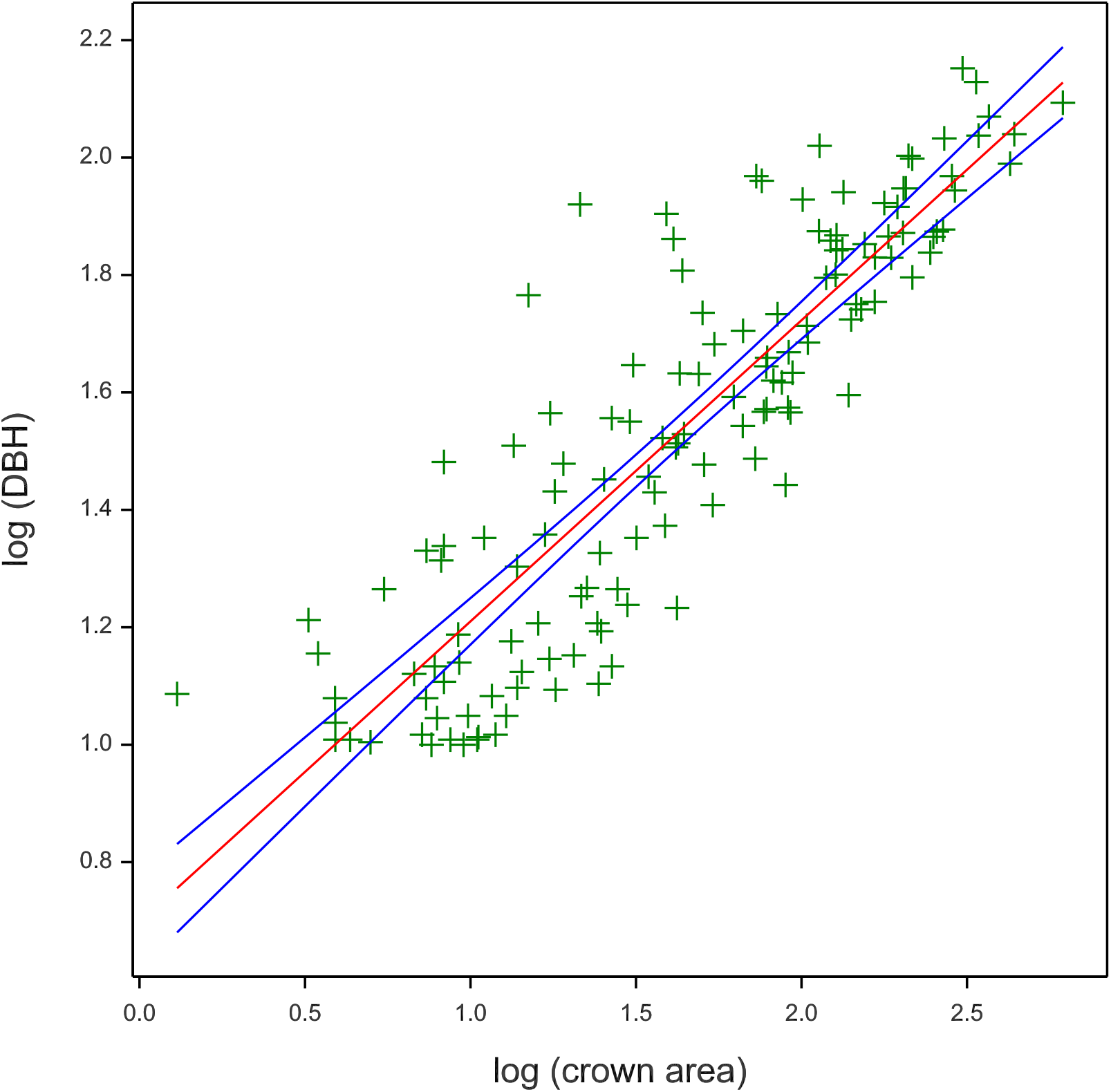
Fitted regression of individual tree stem diameter (DBH, cm) as predicted by crown area (m^2^) for the *T. korupensis* population in the P-plot, using all available data (i.e., including stems with crown area ≤ 75 m^2^). Blue lines show the 95% confidence limits.

**Table S1.** Counts, density, and frequency of newly established seedlings of *T. korupensis* (Tk) [in bold type], *T. bifoliolata* (Tb), and *M. bisulcata* (Mb) sampled in different years in the P-plot at Korup.

|  | No. of plots used | Total area (m<sup>2</sup>) sampled<sup>1</sup> | Tree species | Seedling recruits (N) | Density (N per ha) | Reference source |
| --- | --- | --- | --- | --- | --- | --- |
| Nov. 2010 | 187 | 14 105 | Tb | 780 | 553 | <sup>2</sup>Norghauer and Newbery (2015) |
|  |  |  | Mb | 2118 | 1502 |  |
| Nov. 2008 | 187 | 14 105 | **Tk** | **2415** | **1712** |  |
| Oct. 2007 | 187 | 14 105 | Tb | 1356 | 961 |  |
|  |  |  | Mb | 11 187 | 7931 |  |
| Nov. 1997 | 26<sup>3</sup> | 416 | **Tk** | **201** | **4832** | Newbery et al. (2006b) |
|  |  |  | Tb | 124 | 2981 |  |
|  |  |  | Mb | 200 | 4808 |  |
| Nov. 1995 | 26<sup>3</sup> | 416 | **Tk** | **209** | **5024** | Newbery et al. (2006b) |
|  |  |  | Tb | 360 | 8654 |  |
|  |  |  | Mb | 391 | 9399 |  |
| Nov. 1995 | 91 | 1456 | **Tk** | **135** | **927** | Newbery et al. (1998) |
|  |  |  | Tb | 543 | 3729 |  |
|  |  |  | Mb | 960 | 6593 |  |

<sup>1</sup> This was corrected downward in the 2007, 2008, and 2010 network of systematic sampling, by removing one or more 1/8-wedges of the circular plots, to account for flooded, swampy, or rocky ground.
<sup>2</sup> In that study, seedling recruitment data were used/reported for 111 of the 187 ground plots.
<sup>3</sup> These 26 are subset of the 91 plots established in November 1995 by Newbery et al. (1998).

